# No evidence of induced skin cancer or other skin abnormalities after long term (66 week) chronic exposure to 222-nm far-UVC radiation

**DOI:** 10.1101/2022.03.16.484636

**Authors:** David Welch, Norman J. Kleiman, Peter C. Arden, Christine L. Kuryla, Manuela Buonanno, Brian Ponnaiya, Xuefeng Wu, David J. Brenner

**Affiliations:** Center for Radiological Research, Columbia University Irving Medical Center, New York, NY; Department of Environmental Health Sciences, Mailman School of Public Health, Columbia University Irving Medical Center, New York, NY

## Abstract

Far-UVC radiation, typically defined as 200-235 nm, has similar or greater anti-microbial efficacy compared to conventional 254-nm germicidal radiation. In addition, biophysical considerations of the interaction of far-UVC with tissue, as well as multiple short-term safety studies in animal models and humans, suggest that far-UVC exposure may be safe for skin and eye tissue. Nevertheless, the potential for skin cancer after chronic long-term exposure to far-UVC has not been studied. Here, we assessed far-UVC induced carcinogenic skin changes and other pathological dermal abnormalities in 96 SKH-1 hairless mice of both sexes that were exposed to average daily dorsal skin doses of 396 mJ/cm^2^, 126 mJ/cm^2^ or 56 mJ/cm^2^ of 222 nm far-UVC radiation for 66 weeks, 5 days per week, 8 hours per day, as well as similarly-treated unexposed controls. No evidence for increased skin cancer, abnormal skin growths, or incidental skin pathology findings was observed in the far-UVC exposed mice. In addition, there were no significant changes in morbidity or mortality. The findings from this study support the long-term safety of long-term chronic exposure to far-UVC radiation, and therefore its potential suitability as a practical anti-microbial approach to reduce airborne viral and bacterial loads in occupied indoor settings.

## INTRODUCTION

254 nm UVC radiation emitted from low-pressure mercury lamps is commonly used to disinfect surfaces and room air in hospitals, clean rooms, HVAC supply plenums, and other critical environments (1). However its potential for inducing DNA damage in the skin and eye (2, 3) has limited its direct use in occupied indoor spaces. By contrast, there is now compelling evidence that far-UVC radiation, commonly defined as wavelengths between 200 nm and 235 nm, is likely to be safer for direct human exposure (4-8), and also exhibits similarly or greater antimicrobial activity against both surface and airborne microbes (9-19). The combination of efficacy and safety suggests far-UVC may have broad applicability to provide continuous air disinfection even while humans are present (11, 18).

The biophysical rationale for far-UVC safety is related to the very short penetration depth of far-UVC wavelengths in biological materials (20, 21). Thus, in skin, far-UVC wavelengths are absorbed primarily in the superficial, stratum corneum, which is composed of dead epithelial cells (7, 21, 22) with minimal penetration to the adjacent stratum granulosum, which consists of dead or dying epithelial cells. Biophysical measurements and theoretical calculations imply that far-UVC cannot penetrate to the stratum basale at the base of the epidermis (4, 21), which contains live squamous and basal cells and melanocytes. Damage to cells in the stratum basale is associated with long-term, adverse dermal health effects, including skin cancer (23-25). Recent studies of far-UVC induced DNA photodamage in an in-vitro human 3D skin model support the assertion of a lack of damage to cells in the stratum basale (5, 26).

Correspondingly, in the eye, due to its very short penetration depth, far-UVC wavelengths are absorbed primarily in the tear film, a 3-6 µm acellular protective layer that lubricates and protects the underlying 5-6 layer corneal epithelium (27, 28). Current research suggests that acute ocular health hazards after exposure to even very high exposures of far-UVC may be minimal (29-31).

While theoretical considerations, as well as multiple short-term safety studies in animal models and humans, suggest that far-UVC exposure may be safe for skin and eye tissue, to date there have been very limited safety studies associated with prolonged chronic exposures to far-UVC radiation. Narita, et al. (32), compared far-UVC and 254 nm DNA photodamage in hairless mice exposed to an acute exposure of 450 mJ/cm^2^ for 10 days, but without any analysis of carcinogenic potential due to the limited duration of the experiment. Yamano et al. (33) exposed wild-type and photosensitive *Xpa*-knockout mouse strains biweekly for 10 weeks to acute far-UVC exposures of 500 mJ/cm^2^ or 100 mJ/ cm^2^ respectively. No skin cancers were noted when histological tissue was examined 15 weeks later. While these studies were critical first steps, no study has used the more realistic, prolonged far-UVC exposure scenarios that would be typical for real-life installations intended to minimize disease transmission, where chronic, daily exposures may be delivered over long periods of time.

In the current work, far-UVC induced skin damage was measured in hairless SKH-1 mice exposed to three different irradiances of 222 nm filtered far-UVC radiation for 66 weeks (15.2 months) for 5 days per week and 8 hrs per day. This approach to long-term chronic exposures permits more useful estimations of potential adverse skin health risks arising from the projected use of far-UVC in real-world scenarios to reduce airborne respiratory virus loads.

Hairless SKH-1 albino mice, which were also used in our earlier acute safety studies (10, 34), offer several several advantages for the current study (24, 35). First, the use of a hairless mouse strain eliminates the need for periodically shaving the mouse hair during prolonged chronic exposures. Second, SKH-1 mice are susceptible to UV-induced skin cancer (36-40) and develop lesions resembling human tumors (24, 25, 35, 41). Third, typical stratum corneum thicknesses in mice skin are somewhat less than in humans (5.8±0.3 µm *vs*. 16.8±0.7 µm (42)), so experiments in mice would be expected to be a conservative model for far-UVC effects on human skin. Finally, UVB and UVC (254 nm) induced tumor formation in the hairless mouse has been well-characterized (37-40). These mice develop multiple independent skin tumors, typically from 4 to 10 mm diameter, with characteristic morphology (24). Histologically, these tumors begin as foci of epithelial hyperplasia, progress to papillomas, and ultimately form spindle cell and squamous cell carcinomas, the most common types of UV-associated human skin cancer (43, 44). Such tumors are rarely lethal, and their slow and characteristic progression allows them to be followed non-invasively for many weeks or months.

## MATERIALS AND METHODS

### Animal use and far-UVC exposures

48 male and 48 female, two-month old hairless albino mice (SKH1-Elite Mouse 477; Charles River Labs, Wilmington, MA) were divided into eight groups of 12 mice by sex. Each group was exposed to one of four different 8-hour far-UVC daily doses of nominally 0, 50 mJ/cm^2^, 125 mJ/cm^2^, or 400 mJ/cm^2^ over an eight hour period each weekday (noon to 8 pm) for 66 weeks. These doses were chosen to be higher than the current ICNIRP (International Commission of Non-Ionizing Radiation Protection) Exposure Limit (EL) of 23 mJ/cm^2^ for 222 nm radiation per 8-hour exposure (45), and were motivated by the then-proposed (now adopted (46)) new recommended 8-hour daily Threshold Limit Values (TLV) for 222 nm radiation from the ACGIH (American Conference of Governmental and Industrial Hygienists) of 160 mJ/cm^2^ for the eyes and 480 mJ/cm^2^ for skin (46, 47).

Groups of 12 mice of a single sex were housed together in 8 custom-built 35 × 35 cm acrylic cages fitted with a 79% open area metal mesh top and side ports for food and water to allow *ad libitum* access to each (Fig. 1). Each cage contained wood chip bedding and shredded paper for nesting. Every mouse was identified with a unique radio-frequency identification (RFID) tag (Allflex Electronic Small Animal Identification System, Plano TX) subcutaneously implanted in the scruff of its neck.

**Figure 1.**
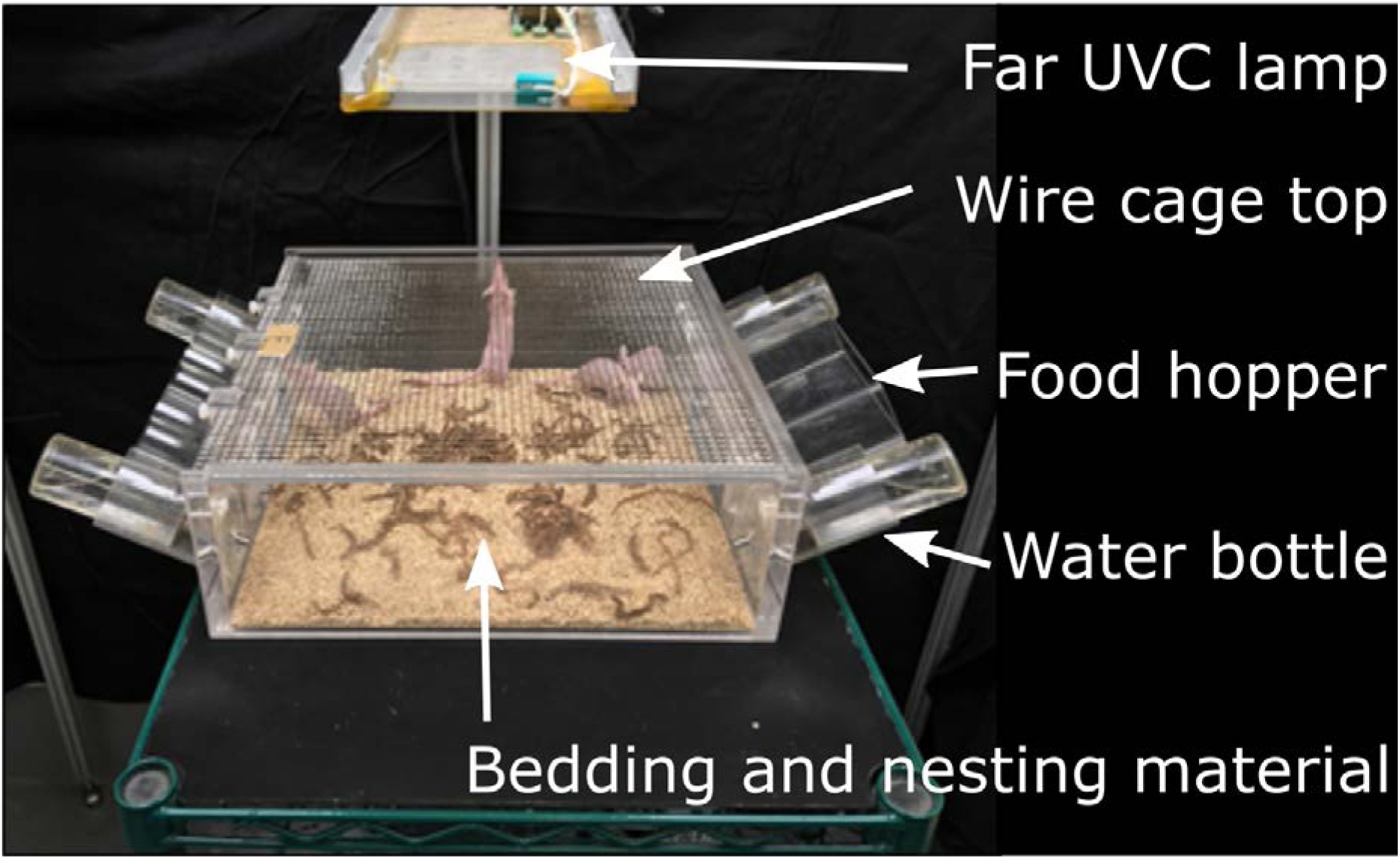
The design of the cage and position of the overhead far-UVC lamp.

The far-UVC sources used in this study were flat, KrCl* excimer microplasma lamps (Eden Park Illumination, Eden Park, IL). KrCl* excimers emit far-UVC radiation with a characteristic 222 nm peak. Custom fabricated optical filters were installed beneath the lamps panels to limit off-peak emissions. For a given dose, each cage of male and female mice was positioned below one of a set of two filtered far-UVC lamps (Fig. 2) whose height was adjusted to provide nominal target irradiances of 400 mJ/cm^2^ per 8-h day (high dose group), 125 mJ/cm^2^ per 8-h day (medium dose group), and 50 mJ/cm^2^ per 8-h day (lowest dose group). In order to average variations in irradiance, the positions of the two cages within a given nominal dose were switched with each other during weekly cage cleanings.

**Figure 2.**
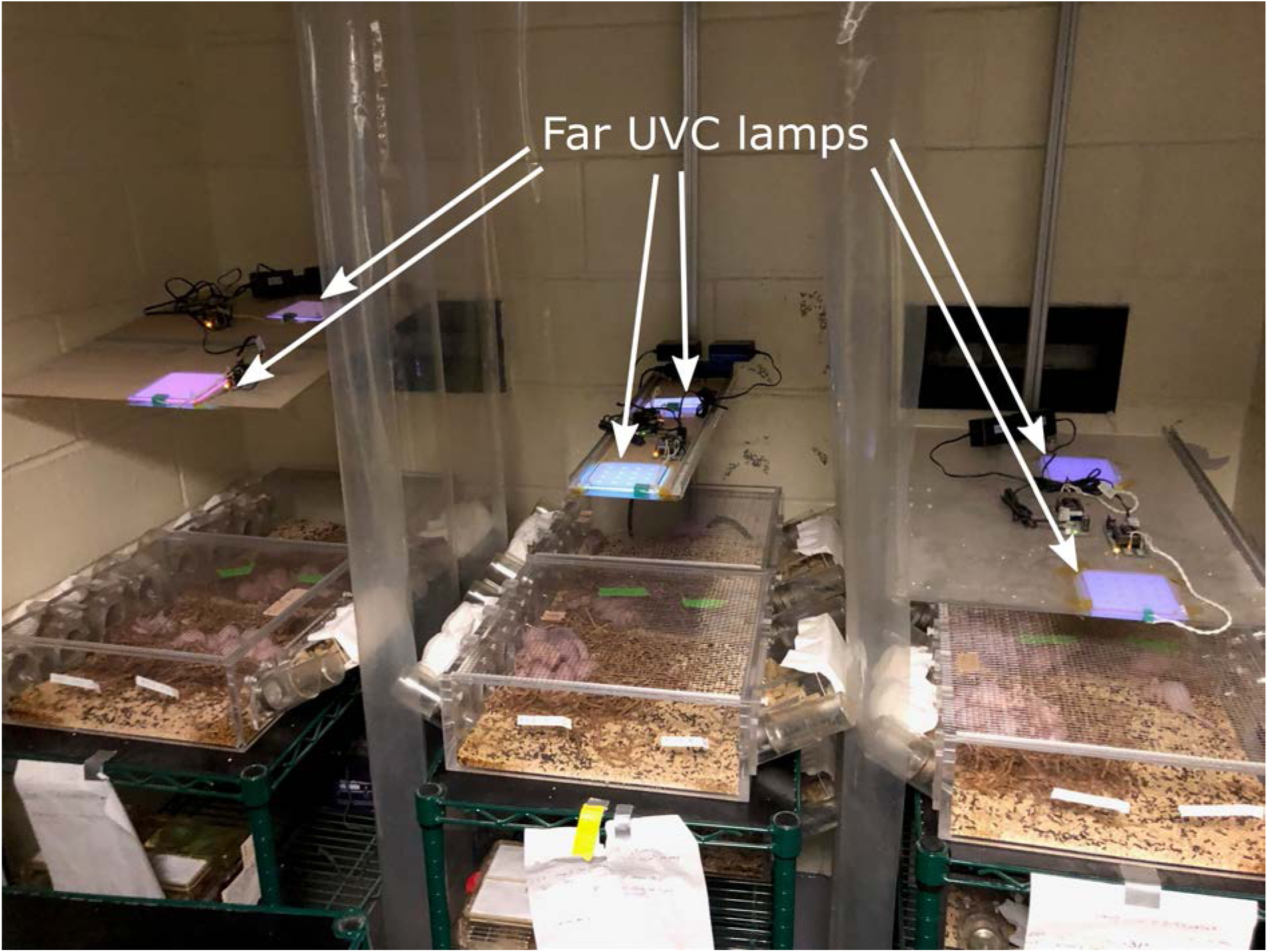
Arrangement of cages and overhead far-UVC lamps. Two far-UVC lamps for each dose were mounted on a vertical rail to allow changes in height above the cage. Higher intensity lamps were used with the high dose positions (leftmost set of cages) and therefore lamps were positioned higher for that position than for the middle dose cohort (middle set of cages) or the low dose cohort (rightmost set of cages). Plastic curtains between cages prevented stray exposures from adjacent lamps. Unexposed control mice were kept in identical cages below the irradiated cages, shielded from far-UVC exposure by opaque shelving.

The cages of the unexposed male or female mice were housed in the same room but were completely shielded from far-UVC exposure. In addition a UVC-opaque plastic curtain was installed between each of the three exposure groups to eliminate stray irradiance from adjacent lamps (Fig. 2).

During weekly bedding changes, each mouse was visually examined for dermal changes as well as for overall health and well-being. After 66 weeks of exposure, the mice were euthanized by cervical dislocation subsequent to ketamine-xylazine anesthesia and abnormal growths or skin lesions surgically removed and immediately placed in 10% formalin for 24 hrs and then washed three times over two days with 70% ethanol prior to paraffin embedding and sectioning.

All procedures and animal husbandry were conducted in accord with accepted ethical and humane practices and carried out in accordance with federal guidelines and an approved Columbia University Institutional Animal Care and Use Committee (IACUC) protocol.

### Far-UVC optical power measurements

Irradiance was measured at the cage floor in a central position using two different optical power meters: a Newport Optical 843-R power meter with an 818-UV/DB sensor (Newport, Irvine, CA) and a Hamamatsu C9536 UV power meter with an H9535-222 sensor (Hamamatsu Corporation, Bridgewater, NJ). The sensitivity of both meters was calibrated throughout the UVC range, and each produced similar measurements of a filtered KrCl lamp with 222 nm peak emissions. Lamp output was measured regularly throughout the 66-week duration of the experiment, and the position of lamps was adjusted to maintain target irradiance values as necessary.

### Far-UVC spectral characterization

Spectral analysis of the filtered KrCl lamps to verify the reduction of spectral emissions of wavelengths other than the 222 nm peak was performed using a Gigahertz Optik BTS2048-UV light meter (Gigahertz-Optik Inc, Amesbury, MA). This spectroradiometer is equipped with cosine-corrected diffusing input optics and has adequate sensitivity and resolution throughout the UV range to enable precise wavelength measurements. The spectral irradiance plot of the filtered lamps is shown in Fig. 3. The six different spectra include measurements at the highest dose (Front and Rear Left), medium dose (Front and Rear Middle), and lowest dose (Front and Rear Right). All spectral measurements were recorded at a distance of 50 mm from the filter. The lower limit of the spectrometer sensitivity is evident with the gaps in measurements and higher levels of noise. The limited spectral contributions outside of the 222 nm peak is indicative of the optical filtration of the typical KrCl spectrum. Differences in the primary 222 nm peak heights are due to variations in lamp output power.

**Figure 3.**
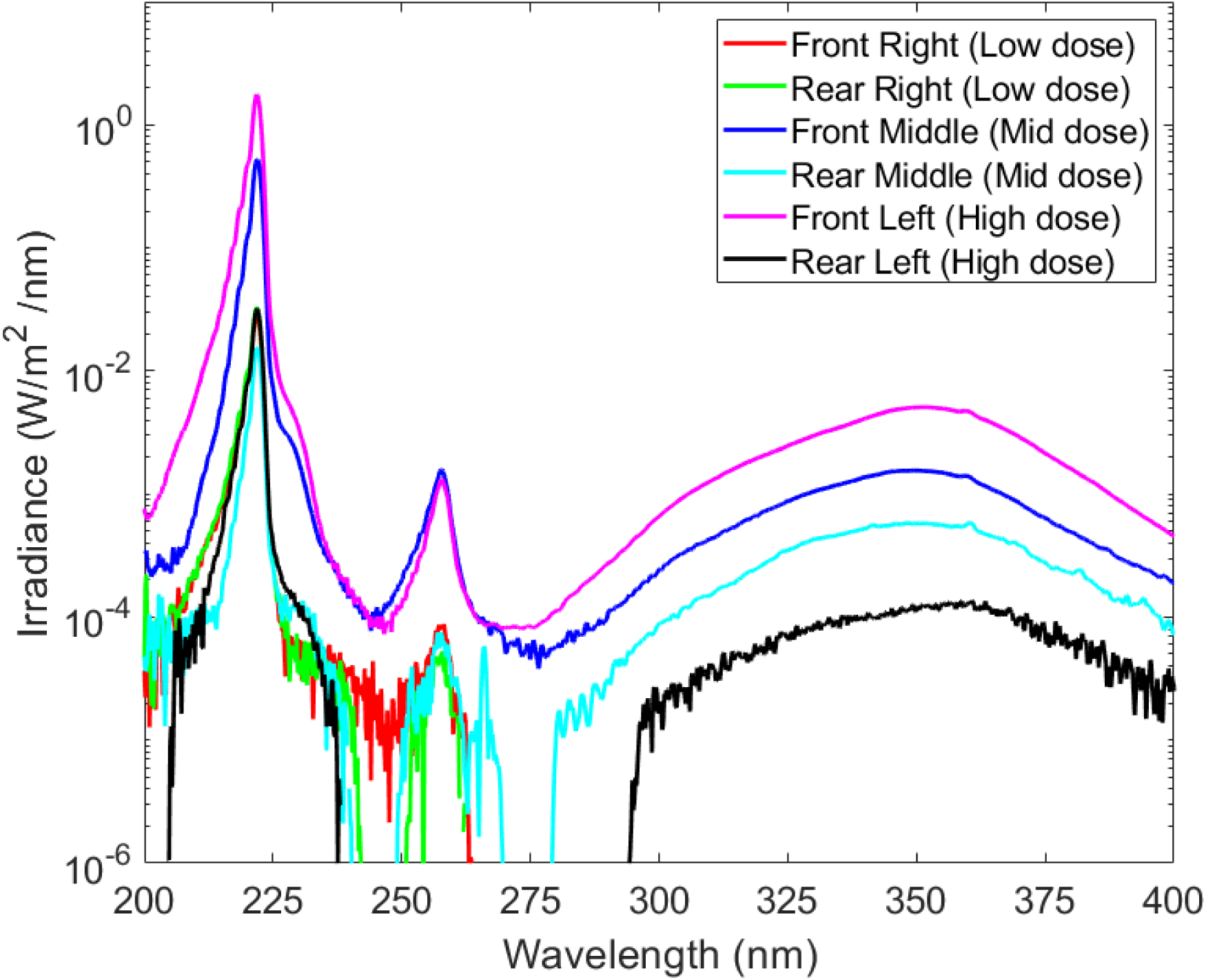
The spectral irradiance measured for the six different exposure positions. All measurements indicate filtering of much of the longer wavelength emissions inherent to a KrCl lamp. Measurements were recorded 50 mm from the filter position.

### Far-UVC dosimetry

Far-UVC dosimetry was performed using calibrated radiation sensitive film (OrthoChromic Film OC-1 (Orthochrome Inc., Hillsborough, NJ)). We have previously demonstrated the utility of OC-1 film for UVC dosimetry (48). This flexible film is 155 μm thick, consisting of a 30 μm active coating on a 125 μm white polyester base. The active region of the film was oriented towards the UV source during measurements since the polyester layer is opaque. The radiant exposure dose received by each film was determined by scanning each film and analyzing using a 222-nm-specific dose calibration curve as described previously (48).

The OC1 films were placed at various locations within each cage to measure the radiant exposure across the cage area. In a separate study (with different mice) to determine actual skin doses to individual mice, 5 × 10 mm strips of OC-1 film were temporarily glued to the back of each mouse using veterinary surgical adhesive (GLU-CA1999-BUT, Covetrus, Dublin, OH). Up to twelve mice were used for each measurement, and the dosimetry cage was positioned under each of the six lamp locations used in the study (Fig. 2). An example of the experimental setup is shown in Fig. 4. After approximately three hours exposure, the film was gently peeled off from each mouse, the film color density was quantified using a flatbed scanner, and the cumulative dose during the exposure time was estimated as previously described (48). A three-hour exposure was chosen as a balance between a sufficient length of exposure and the propensity for the film to either fall off or be removed by the mice during grooming. Only films that remained attached to the backs of mice throughout the entire ∼3 hour exposure time were included in the dose assessment. The average irradiance upon the film for the exposure time was used to extrapolate the 8-hour skin dose. This method of film dosimetry was performed twice at each of the 6 cage positions.

**Figure 4.**
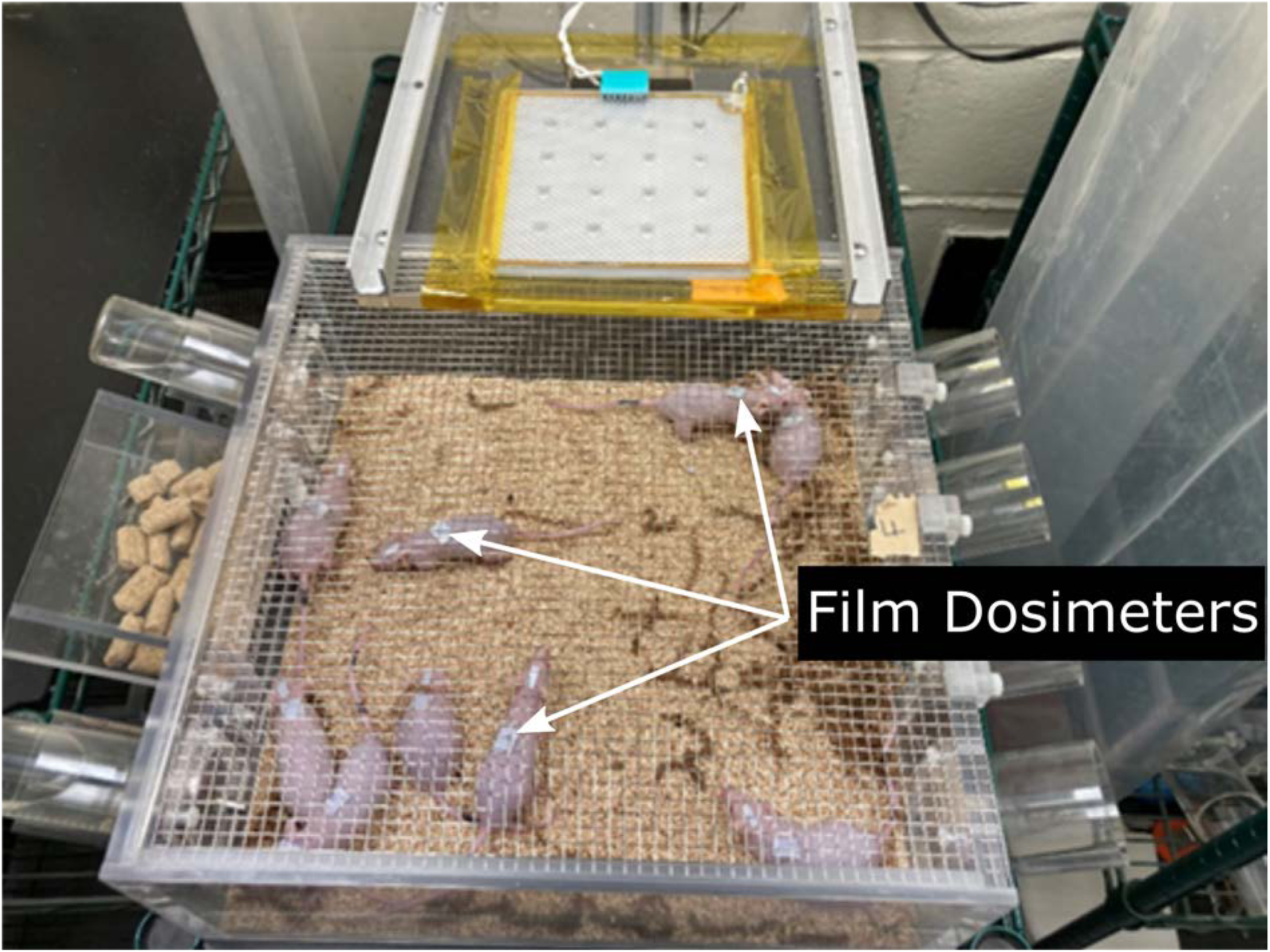
A cage of mice with film dosimeters (light blue) affixed to their backs for individualized dosimetry measurements.

### Statistical analysis

Comparisons between the results in the four dose groups, as well as sex comparisons, were performed using Fisher’s exact test. Survival curves were analyzed using Kaplan-Meier methodology and compared using log rank tests, and mass growth curves were compared using the CGGC (Compare Groups of Growth Curves) permutation test tool (49).

## RESULTS AND DISCUSSION

### Experimental UV Dosimetry and Characterization

Irradiance values measured using an optical power meter centered on the floor of each cage were used to determine the lamp positions for each of the target doses, and a film dosimetry approach was used to examine irradiance values and the extrapolated 8-hour radiant exposure dose within the cage area with high spatial resolution. Film dosimetry measurements were made across the area of the cage at a 20 mm height above the cage floor to approximate the plane of the height of the back of a mouse. A two-dimensional map of the irradiance across each cage was constructed using the results, and the 8-hour extrapolated radiant exposure dose is illustrated in Fig. 5. At all doses, this analysis revealed that the irradiance was higher in the center of the cage due to the emission pattern of the lamp.

**Figure 5.**
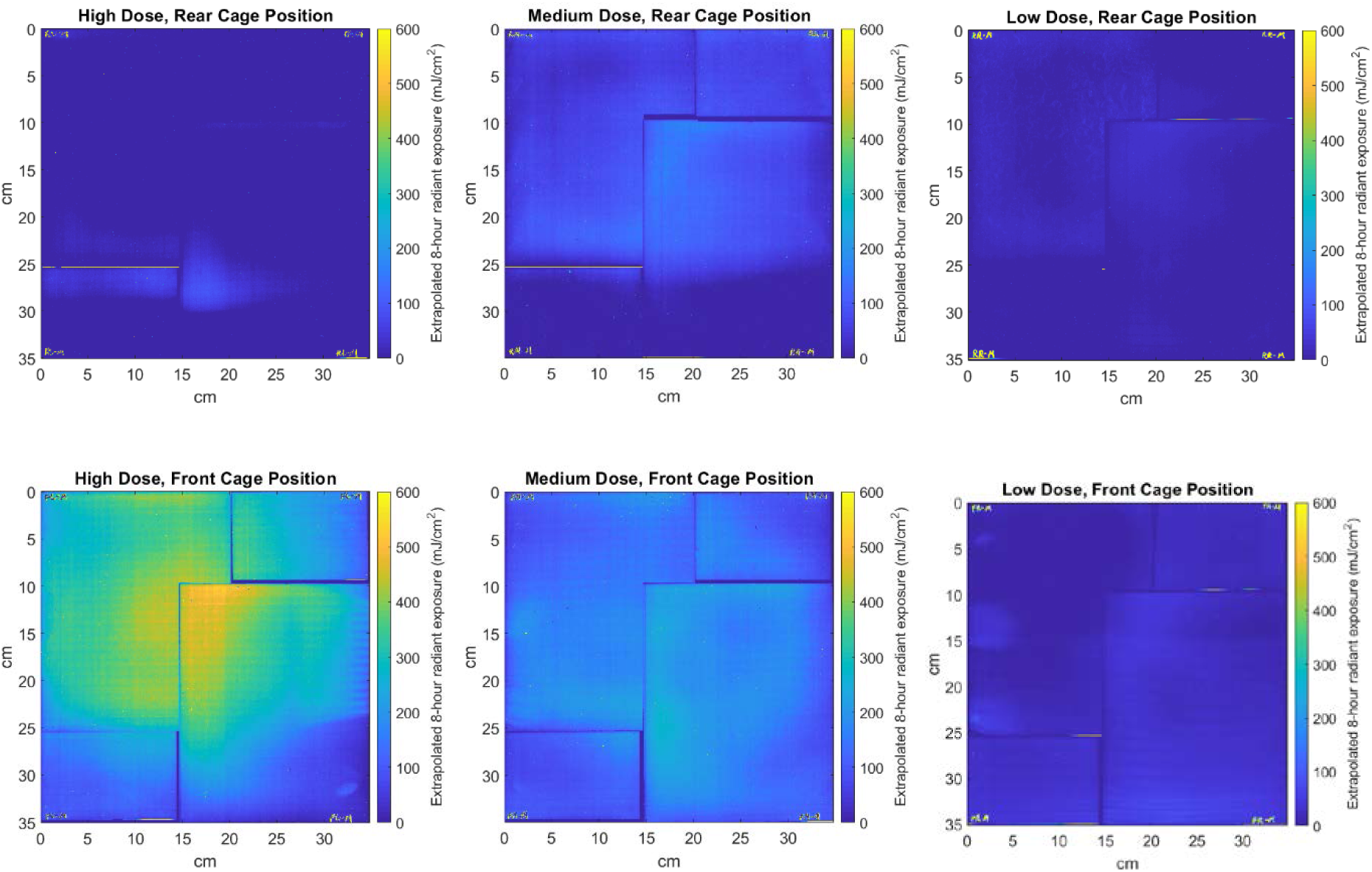
The measured exposure field within the cage areas is shown for each of the six cage positions, with the extrapolated 8-hour daily radiant exposure dose indicated by the color map. The leftmost column shows the two high-dose positions, the middle column shows the medium dose positions, and the right column shows the low dose positions. Multiple sheets of film were required to span the entire cage area, and the small spaces between sheets are visible in the dose map.

Observations over the course of the 66-week exposure revealed multiple instances of mice huddled together in the cage corner, in an upright position exploring the cage walls or eating or drinking, and occasionally in inverted positions in which a mouse was hanging upside down from the wire mesh on the top of the cage. The variation in irradiance across the cage area, along with the dynamic position of each mouse within the cage over the course of the exposure time, necessitated the use of a novel individualized in-vivo dosimetry method. As described in “Methods”, this was accomplished using small pieces of dosimetric film attached to the backs of individual mice which were freely moving within the cage over a test period (these were different mice, “dosimetry mice”, than those used for the full 66 week study). Both variations in the mouse position and variations in the irradiance within the cage were accounted for using this in-vivo dosimetric method. The average extrapolated 8-hour exposure doses to the backs of the mice for both cage positions are given in Table 1. As the cages were interchanged weekly between their two positions, the average values for the two cage positions for each target dose was considered the daily exposure dose for the mice for the 66-week exposure period.

**Table 1.** Calculated average 8-hour dose to the backs of each mouse housed in each of the six cage positions. The mean and standard deviation at each cage position was calculated from the number of films which remained affixed throughout the dosimetry testing period of ∼3 hours. Because the cages were rotated between the front and rear positions on a weekly basis, the average of the two cage positions was taken as the average dose over the total exposure time.

|  | Low dose cages | Medium dose cages | High dose cages |
| --- | --- | --- | --- |
| **Rear cage position** | $48 \pm 7.5 \text{ mJ/cm}^2$ (n=17) | $67 \pm 25 \text{ mJ/cm}^2$ (n=23) | $79 \pm 8.8 \text{ mJ/cm}^2$ (n=19) |
| **Front cage position** | $65 \pm 9.3 \text{ mJ/cm}^2$ (n=18) | $185 \pm 27 \text{ mJ/cm}^2$ (n=22) | $712 \pm 120 \text{ mJ/cm}^2$ (n=22) |
| **Average dose** | **$56 \text{ mJ/cm}^2$** | **$126 \text{ mJ/cm}^2$** | **$396 \text{ mJ/cm}^2$** |

As shown in Table 1 and Fig. 5, for each dose group there were clear differences between the doses for the front and the rear cage positions. However, as each cage was switched between the front and rear location on a weekly basis, it is reasonable to estimate the average 8-hour daily dose over the 66-week exposure period as the average of the two cage positions for each exposure dose.

### Skin observations and histological measurements

Over the course of the 66-week exposure period, each mouse was comprehensively examined for visual abnormalities and any abnormal findings noted and recorded. If observed, any abnormalities were monitored non-invasively at subsequent examinations. Mice weights were recorded at ten of these examinations.

After 66 weeks, all surviving animals were euthanized and abnormal growths or tissue abnormalities excised and sent for pathological examination. After examination by a board certified veterinary pathologist, none of the samples exhibited any evidence of neoplasia or any other pre-cancerous change. No squamous cell carcinomas, pre-malignant papillomas, or malignant microinvasive squamous cell carcinomas were observed. The most common findings in the control and irradiated mice were multiple dermal cysts, utriculi, and/or hyperplastic sebaceous glands, all of which are consistent with those described in aging mice in general and, in particular, in the SKH-1 mouse strain (35, 50).

These histological results are summarized in Table 2. As mentioned above no skin cancers were seen in any of the groups. The number of mice exhibiting non-cancerous skin abnormalities in each of the four dose groups (control, low, medium, high) were compared and the combined results from the three irradiated groups were also compared with the control group results. For all these pairwise comparisons, performed with Fisher’s exact test, we could not reject (p>0.05, 95% confidence) the null hypothesis that the histological results were not significantly different between the different does groups, and were not significantly different between the two sexes.

**Table 2.** Skin changes and survival across groups. Pathological evidence of skin carcinogenesis was operationalized as neoplasia or any other pre-cancerous change, squamous cell carcinomas, pre-malignant papillomas, or malignant microinvasive squamous cell carcinomas (none were found).

| Dose Group↓ | Sex | Number of mice at 0 wks | Number of live mice at 66 wks | Pathological evidence of skin cancer | Incidental pathology findings |
| --- | --- | --- | --- | --- | --- |
| Control | F | 12 | 7 | 0 | 2 |
|  | M | 12 | 10 | 0 | 5 |
| Low Dose | F | 12 | 8 | 0 | 1 |
|  | M | 12 | 9 | 0 | 3 |
| Medium Dose | F | 12 | 9 | 0 | 5 |
|  | M | 12 | 9 | 0 | 4 |
| High Dose | F | 12 | 10 | 0 | 3 |
|  | M | 12 | 10 | 0 | 4 |
| Total |  | 96 (48 F, 48 M) | 72 (34 F, 38 M) | 0 | 27 (11 F, 16 M) |

### Weight changes and mortality

An indicator of potential carcinogenesis in mice, often before detection of overt tumor formation, is abnormal weight change (51). Thus all animals were weighed on a regular basis, and the time time-dependent weight changes are shown in Fig. 6. The results for the four exposure groups were compared using the CGGC test tool (49) and no statistical difference between the weight-change curves between the four exposure groups were seen, both for both sexes combined (Fig. 6) or when further stratified by sex.

**Figure 6.**
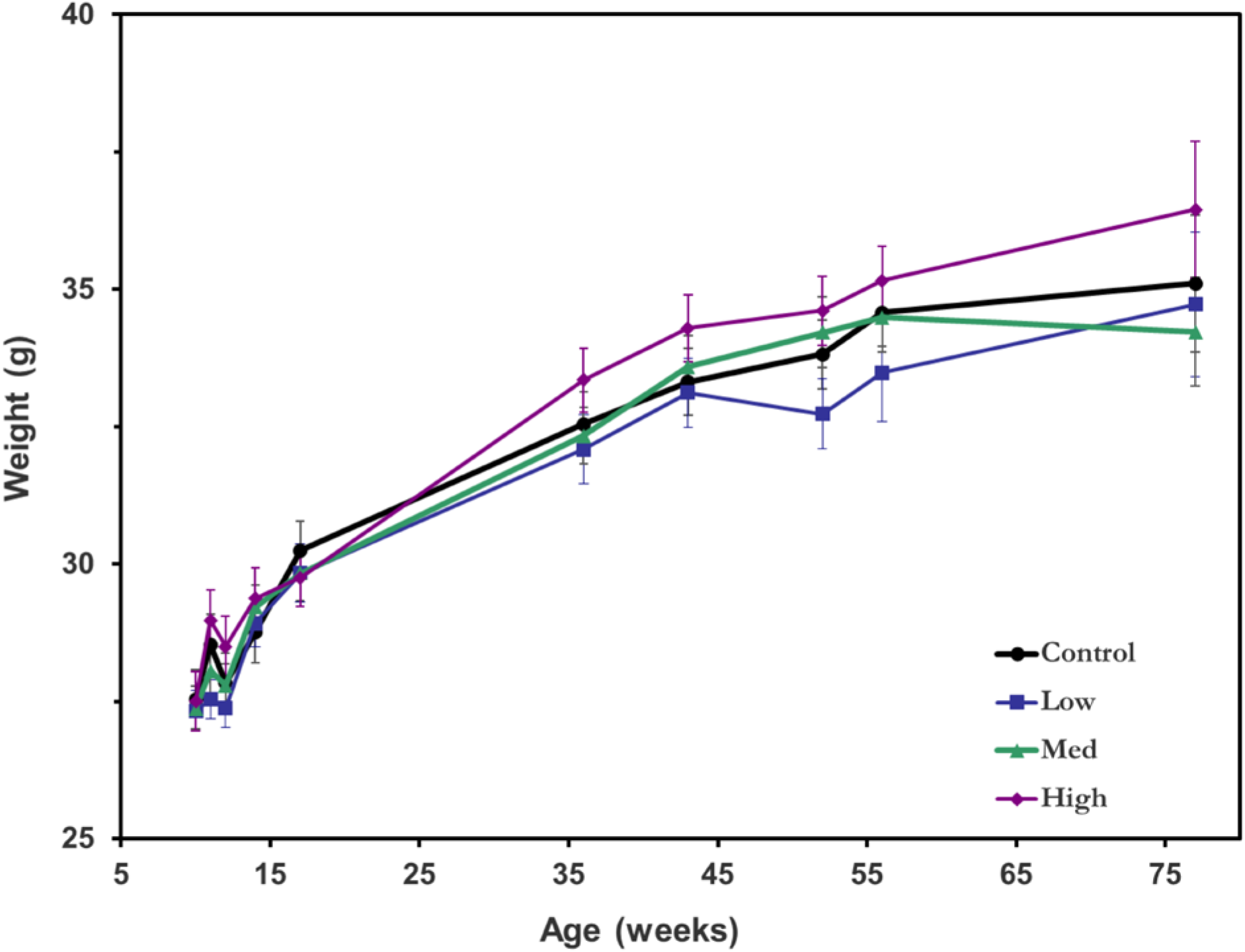
Average mouse weight over the irradiation period in each of the four exposure groups. Error bars indicate standard errors.

We also analyzed excess mortality and Fig. 7 shows Kaplan-Meier survival curves for each of the three exposure groups and the overall unexposed controls. Using standard log-rank tests, no statistical difference in survival was observed between any of the four groups (p>0.05). No significant difference was obtained when the results were further stratified by sex.

**Figure 7.**
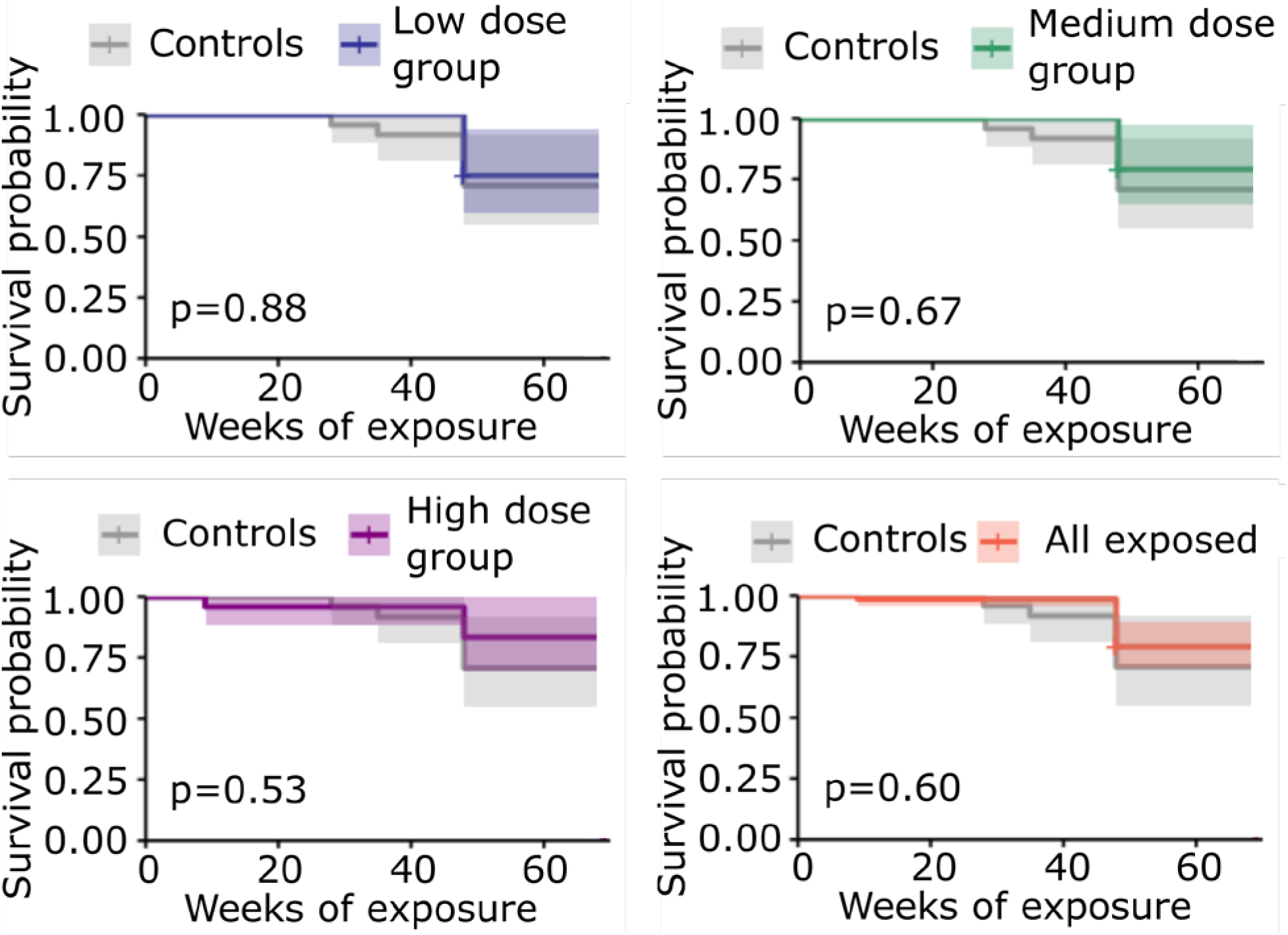
Kaplan-Meier plots and p-values for each exposure dose group (low, medium, high, or all groups combined) compared to the unexposed control group. Shading represents 95% confidence intervals.

Mortality was also compared by analyzing the numbers of mice that survived in each of the four exposure groups at the end of the 66-week study (Fig. 8), with confidence intervals calculated using standard methods for binomial proportions (52). Based on Fisher’s exact tests, no statistical difference in mortality between any of the exposure groups was detected, whether analyzed separately by sex or with sexes combined.

**Figure 8.**
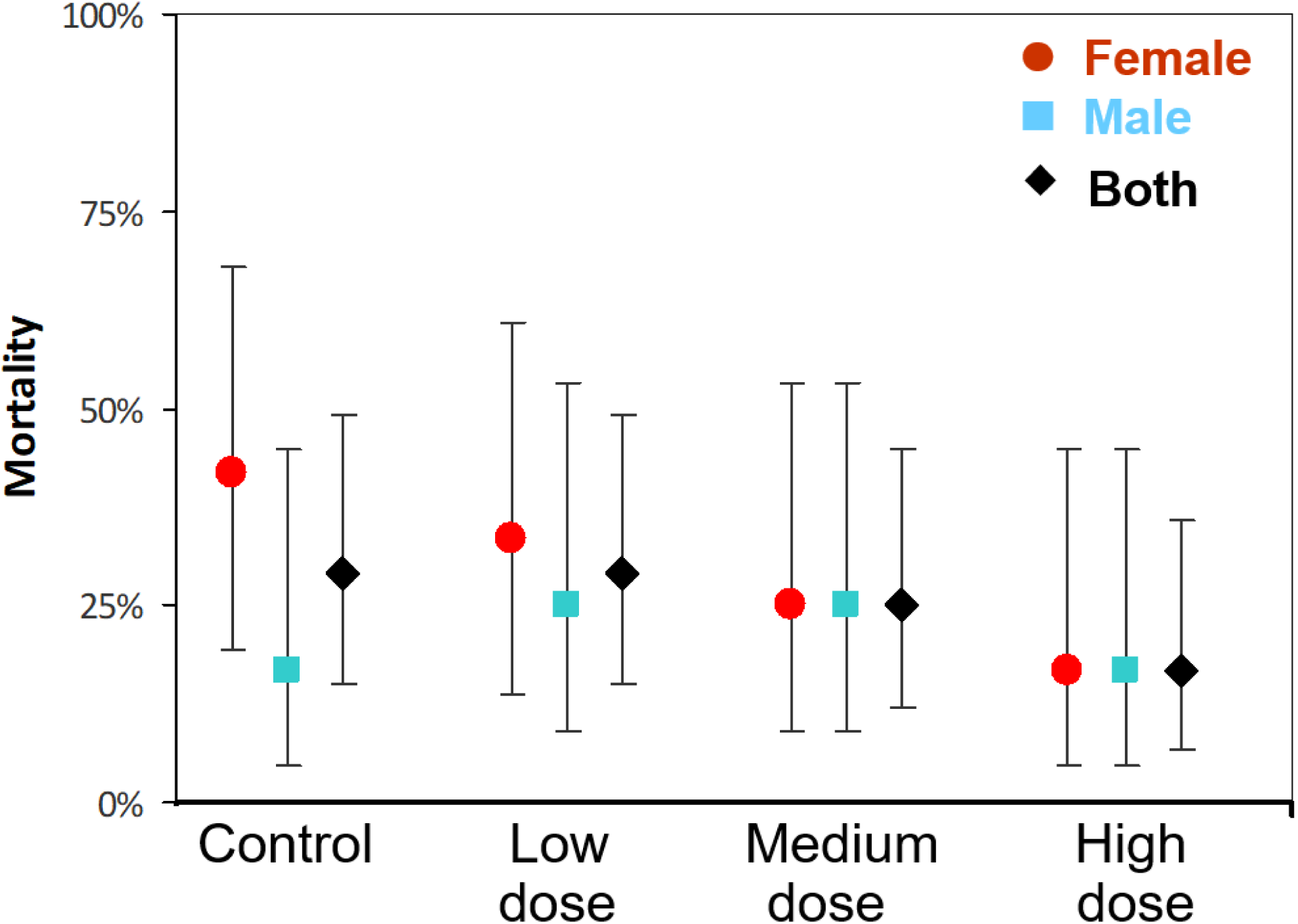
Mortality (Percent of mice that did not survive to the end of the 66 week study) by exposure cohort (unexposed control, low, medium, and high dose) separated by sex (male-blue, female-red) and combined for both sexes (black). 95% confidence intervals are shown.

## CONCLUSIONS

In summary, the findings of this study indicate that chronic exposure of SKH-1 hairless mice to far-UVC 222 nm radiation for 66 weeks (15.2 months) at mean daily 8-hour skin doses of up to 400 mJ/cm^2^ did not result in any evidence for induced skin cancer in SKH-1 hairless mice. In addition, we found no evidence of far-UVC related increases in non-cancerous skin lesions, unusual weight loss, or excess mortality. The SKH-1 hairless mouse is a standard model which has been used to quantify the significant yields of skin cancers and other skin lesions induced by prolonged exposure to UVB radiation (39) and prolonged exposure to conventional germicidal UVC (254 nm) radiation (38).

This is the first study to estimate skin cancer risk from chronic, long-term far-UVC exposure and the negative results here offer insight into the safety of human skin after prolonged 222 nm exposure. The negative findings reported here are pertinent to the potential use of far-UVC technology to control airborne microbe transmission in occupied indoor spaces.

## Acknowledgments

This work was supported by the Shostack Foundation, NIH grant 5R42AI125006, and the Boeing Company. We thank Dr. Gerhard Randers-Pehrson for his conceptual insights.

